# INDITEK-2.0: A Bayesian inverse eco-evolutionary modelling framework for reconstructing Phanerozoic biodiversity

**DOI:** 10.1101/2025.08.22.671786

**Authors:** Pedro Cermeno, Carmen García-Comas, Gloria Herrero Gascón, Michael J. Benton, Thomas Bodin, Khushboo Gurung, Benjamin Mills, Alexandre Pohl

## Abstract

1. Mechanistic eco-evolutionary models are powerful tools for understanding how life diversified over geological timescales, yet estimating their parameters remains a key challenge. The high computational requirements of these models make probabilistic inversion methods unfeasible.
2. We address this challenge with INDITEK-2.0, a new, cost-efficient eco-evolutionary model that integrates a Bayesian inverse modelling framework for probabilistic estimation of parameters.
3. To validate this framework, we conducted a proof-of-concept study. We first generated synthetic biodiversity data for marine invertebrates using INDITEK-2.0 with known parameter values. To mimic field data, some realistic random Gaussian errors were added to the data. We then used our Bayesian inverse modelling framework to recover these original (true) parameter values from the dataset of current biodiversity distributions after 500 million years of diversification. Our solution is a probability distribution on the parameter space, and we show that the method successfully recovered the target parameter values with a relatively low degree of uncertainity.
4. INDITEK-2.0’s ability to probabilistically infer underlying eco-evolutionary processes provides a new modelling tool for reconstructing the evolutionary history of biodiversity across Earth’s ecosystems and taxonomic groups.

## 1. Introduction

Mechanistic modelling is emerging as a powerful complementary tool to the fossil record and molecular phylogenetics for reconstructing biodiversity history (Beaugrand, 2023; Hagen, 2023; Lovelace et al., 2020; Rangel et al., 2018). By simulating eco-evolutionary processes under dynamic environmental conditions, mechanistic eco-evolutionary models offer a unique opportunity to explore the drivers and patterns of taxonomic diversification, bridging gaps and testing hypotheses that are difficult to address with traditional methods alone. They notably permit to go beyond the observation of temporal correlations between environmental change and biodiversity dynamics and, through numerical experiments, to address causality and identify the mechanisms underlying observed changes. Mechanistic eco-evolutionary models have been developed to date at two levels of biological organization, at the individual level and the population level. Individual-based eco-evolutionary models provide a framework for simulating complex ecological and evolutionary processes, with trait variation and demographic stochasticity modelled at the level of individual organisms, each represented as a unique entity with its own set of traits (e.g., size, age, genotype, phenotype) (Garwood et al., 2019; Malchow et al., 2021; Xuereb et al., 2021). By incorporating genetic and phenotypic trait variation, individual-based eco-evolutionary models can predict how populations respond to environmental changes, including habitat fragmentation, climate change, or the introduction of invasive species. Population-based models, on the other hand, abstract the details of individual organisms into population or species-level traits that influence species’ survival, reproduction, range-shifts, and interaction with the environment and other organisms (Beaugrand, 2023; Hagen et al., 2021; Rangel et al., 2018). The distribution and frequency of these traits within the population can change over time through natural selection, genetic drift, and dispersal, reflecting the evolutionary processes of speciation, extirpation, and extinction at play.

To capture the details of eco-evolutionary processes, individual- and population-based eco-evolutionary models must operate at sufficiently high spatial and temporal resolutions. High-resolution modelling represents habitat heterogeneity and species variability more realistically than coarser models. However, this level of detail creates computational challenges when reconstructing evolutionary histories over vast spatial and temporal scales. This is a problem for both the Earth system models that provide palaeoenvironmental conditions and the eco-evolutionary models themselves. Likewise, in a mechanistic eco-evolutionary model, each discrete entity or state variable, whether an individual organism or a population of organisms, is governed by its own set of differential equations that describe its state changes over time. These equations account for growth, reproduction, death, migration, interactions with other entities and the environment, genetic drift, heritability, speciation, and extinction. The need to solve a separate set of equations for each discrete entity rapidly escalates the computational demands of these models.

Beyond current computational limitations, estimating eco-evolutionary model parameters (evolutionary rates, environmental dependencies, carrying capacities) requires detailed knowledge of the very biodiversity history we aim to reconstruct, which creates a circularity problem. Inverse modelling offers a solution: it compares model simulations with empirical data (e.g. present-day biodiversity, fossil data). By finding the simulation that best fits the empirical data, inverse modelling can effectively estimate the parameters that most likely generated the observed patterns (Tarantola, 2005). Probabilistic inference, particularly Bayesian methods employing Markov chain Monte Carlo (MCMC) algorithms, provides a robust methodological framework to accomplish this. MCMC algorithms explore the multidimensional parameter space by taking random walks guided by the shape of the posterior distribution. By recording the visited locations within the parameter space, MCMC builds a representative sample of the posterior distribution. While effective, this method requires thousands of model simulations for adequate parameter space exploration, adding a significant layer of computational requirements to these already demanding models. Consequently, applying Bayesian inversion to these models requires a trade-off between model complexity and computational feasibility.

To address this trade-off, Cermeno et al. developed INDITEK, a cost-efficient eco-evolutionary model that simplifies mechanisms while still capturing essential broad-scale, macroevolutionary dynamics (Cermeño et al., 2022). Previous models use a bottom-up approach, modelling biodiversity by individually tracking individuals or species. In contrast, INDITEK utilizes a top-down approach, treating biodiversity as an aggregate property at a taxonomic level greater than the species level (e.g., class, phylum). This methodological shift moves the analytical focus from individual- or species-level traits to broader attributes at class or phylum level, making biodiversity itself—rather than individuals or species—the state variable. This strategic approach overcomes major computational hurdles by implicitly modelling the phylogenetic divversification process, allowing the simulation of over 500 million years of global marine invertebrate diversity in under thirty seconds on a standard computer. This article presents INDITEK-2.0, an upgraded version of the INDITEK model. INDITEK-2.0 integrates a Bayesian inverse modelling framework, which enables probabilistic estimation of its parameters. We first conduct a proof-of-concept study to assess the framework’s reliability using synthetic community data. We then evaluate its applicability to macroevolutionary questions, concluding with a discussion of model limitations and future directions.

## 2. INDITEK-2.0 theoretical framework

### 2.1. Parameterizing evolutionary rates

As noted, in our model, biodiversity is an aggregate property of the community implicitly reflecting the cumulative effects of ecological and evolutionary processes at or below the species level. These processes give rise to spatial and temporal differences in diversification dynamics, whose parameterization is critical to reconstructing and understanding the evolutionary history of biodiversity.

The metabolic theory of ecology provides a theoretical basis, with substantial empirical support, for understanding how temperature affects evolutionary rates by altering the energy available for growth, reproduction, and survival (Allen et al., 2002; Brown et al., 2004). A greater energy input to the system (or a higher ambient temperature) implies faster metabolic rates and, consequently, shorter generation times (Rohde, 1992). Higher turnover rates allow for quicker accumulation of genetic changes, potentially increasing the rate of evolution (Lanfear et al., 2010). This means that adaptive traits can become prevalent in shorter time spans, allowing populations to respond to selection pressures more rapidly (**Fig. 1**). The metabolic theory of ecology uses the Boltzmann factor, a concept from statistical thermodynamics, to explain how the rates of biological processes, like metabolism, growth and reproduction accelerate with temperature (Gillooly et al., 2002). Its core idea – that probabilities can exponentially change with energy and temperature – can be applied to parameterize how temperature affects speciation rate through its impacts on biological rates. Alternatively, another quantitative expression for parameterising the sensitivity of metabolic rates to temperature changes is the Q10 coefficient, a dimensionless parameter taken from the field of ecophysiology (Mundim et al., 2020). The simplicity of the Q10 model – offering a numerical value to represent how metabolic rates change at a different rate with every 10°C change – makes it particularly suitable to eco-evolutionary modelling (**Box 1**).

**Fig. 1.**
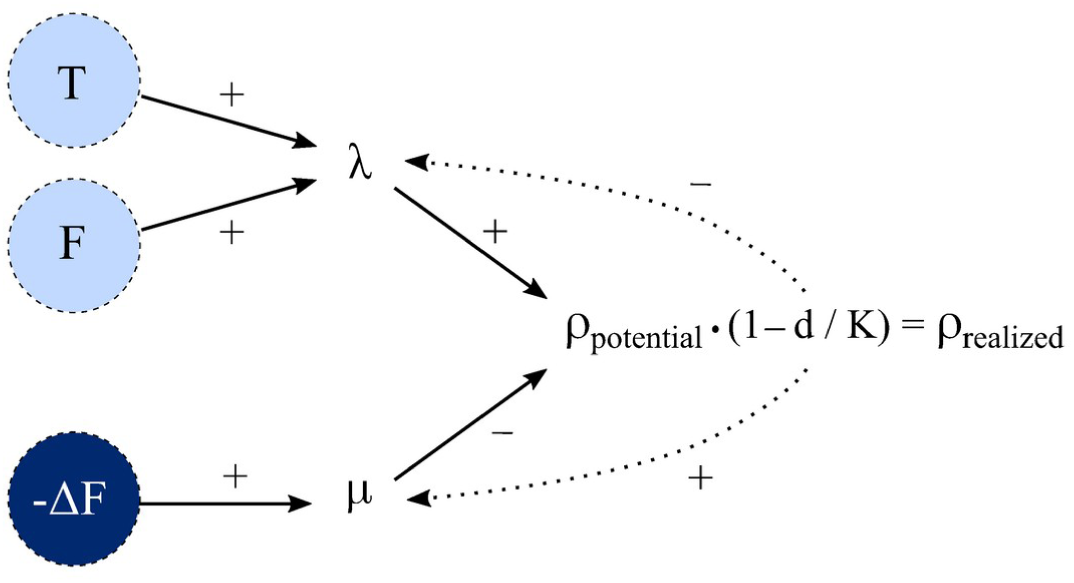
Parameterization of evolutionary rates. Speciation rate (λ) is a positive function of temperature (T), food availability (F), and, in terrestrial systems, water availability (not shown). A reduction in food supply rate (-ΔF) directly lowers the carrying capacity (K), implicitly driving an increase in the extinction rate (μ) in systems near K. The difference between λ and μ determines the net diversification rate, ρ_potential_. The actual, or realized, net diversification rate (ρ_realized_), is calculated by multiplying ρ_potential_ by the diversity-dependent limitation term (1-d/K). Diversity-dependent limitations feedback on evolutionary rates by implicitly decreasing λ, increasing μ, or both as d approaches K (dotted arrows) .

Our parameterization allows species to evolve their thermal tolerance range—the limits that influence their speciation rate—to track the prevailing climate conditions across each geological time interval (**Box 1**). For example, a temperature that is considered an extreme heat event in the Ordovician period might be a normal temperature in the much warmer Eocene epoch. This approach prevents the model from assuming a universal thermal tolerance for organisms across the entire Phanerozoic, which would be biologically unrealistic. It ensures that the influence of temperature on speciation rate is assessed based on deviations from the norm of that time, thus providing a more accurate and biologically meaningful representation of the relationship between temperature and speciation rate.

Food supply also plays a pivotal role in controlling speciation rates by affecting population dynamics and energetics (Bambach, 1993; Martin, 1996; Vermeij, 1977). For example, abundant food resources can support larger populations, which tend to maintain higher genetic diversity (Hague & Routman, 2016). Larger, genetically diverse populations provide a broader repertoire for natural selection to act upon, potentially accelerating the speciation process (**Fig. 1**). Similarly, for terrestrial ecosystems, water availability, primarily through precipitation, is a fundamental control on net primary productivity (NPP), which in turn determines the available food supply. As a key component of photosynthesis, water is a limiting resource in many ecosystems, directly influencing the amount of biomass produced and thus acting as a primary driver of speciation rates. As with temperature, quantifying the combined effect of food and water availability on speciation rate necessitates the use of generalized mathematical models. These models, ranging from simple linear relationships to more complex nonlinear dynamics, provide a quantitative framework for representing the mechanisms that govern this relationship (**Box 1**). For example, a Michaelis-Menten equation provides a useful quantitative model for describing the dependence of speciation rates on both food and water availability. This approach is analogous to how chemical reaction rates depend on substrate concentration in biochemistry. It can capture the threshold-like behaviour of resource-driven productivity, where speciation rates increase rapidly with increasing resources in limited environments but plateau in resource-rich conditions where other factors become limiting.

Parameterizing the rate of extinction is considerably more difficult because there is no clear, well-established theoretical framework that comprehensively explains the mechanistic drivers behind it. Adaptation is a fundamental evolutionary process allowing species to survive in changing environments. However, significant changes in environmental conditions can outpace a species’ ability to adapt, resulting in a mismatch between its physiological requirements and the new environmental conditions (Hallam, 1998; Peters & Foote, 2002; Saupe et al., 2014). When these changes occur rapidly, species may not have enough time to evolve new traits and cope with the new conditions. For example, it has been hypothesized that the latitudinal biodiversity gradient, characterized by the increase in species richness towards the equator, might result from greater environmental instability at higher latitudes (Chown & Gaston, 2000; Willig et al., 2003). This reasoning posits a direct linkage between the rate of environmental change and the rate of species extinction (Kaiho, 2022; Song et al., 2021). This parameterization requires, however, higher spatial and temporal resolution environmental data than is currently available (see ***Section 4. Model limitations and future directions***). While future Earth system model simulations may provide the necessary inputs, here, we assume a constant background extinction rate for the current model.

### 2.2. Diversity dependence of net diversification rate

Simulating the effect of ecological interactions, such as competition or predation on diversification rates requires accounting for species interactions, which is not possible in a model operating above the species level. One way to incorporate the effect of biotic factors without getting into the complexities of species interactions is by using the concept of carrying capacity (K) for species richness (Storch & Okie, 2019). The parameter K is intricately tied to the balance between the available resources in the system and the species that depend on these resources for survival. On evolutionary time scales, the dynamic interplay among resource availability and the strategies species employ to exploit and compete for these resources establishes K, the maximum number of taxonomic units an ecosystem can support (Bambach, 1983; Mondal & Harries, 2016). In the context of evolutionary models, K can be used to approximate the effect of ecological interactions (or biotic factors) on evolutionary rates. As diversity (d) approaches K, the strengthening of ecological interactions slows down the diversification process until equilibrium diversity is reached (Levinton, 1979; MacArthur & Wilson, 1967). These diversity-dependent ecological interactions feedback on evolutionary rates by implicitly decreasing λ, increasing μ, or both (Rabosky, 2009; Rineau et al., 2022) (**Fig. 1**).

In our model, the K of an ecosystem is directly tied to its net primary production (NPP), which we use as a proxy for food supply. We reconstructed marine and terrestrial NPP coupling a long-term carbon cycle model (SCION) (Mills et al., 2021) with an Earth system model of intermediate complexity (cGENIE) (Ridgwell et al., 2007) (see **Supporting information**) and established a global minimum K (K_min_) and maximum K (K_max_) value for each realm (marine and terrestrial). The actual K for a specific location and time was then calculated as a linear function of the NPP at that specific point, with a range that encompassed the values between K_min_ (lowest NPP) and K_max_ (highest NPP) (see **BOX 1** for details). This approach hinges on the idea that NPP sets a fundamental limit on the available chemical energy (or food) to support consumer populations (Cárdenas & Harries, 2010; Martin, 1996). Accordingly, when a decrease in NPP occur due to environmental shifts, the ecosystem’s K is consequently lowered. In response to this resource limitation, biodiversity is reduced mimicking a local extinction event to align with the new, lower K (**Fig. 1**). In this way, and according to theory, our model links biodiversity directly to the dynamics of energy input to ecosystems (Coelho et al., 2025; Hawkins et al., 2003; Wright, 1983). Ultimately, K at a given location and time is determined by both NPP and the values of K_min_ and K_max_. Yet, these latter values should not be static as in the current model; rather, they are expected to change over geological time, driven by evolutionary innovation (see *4. Caveats and future directions*).

#### Box 1 INDITEK-2.0’s model parameterization

To reconstruct biodiversity, we employ a logistic differential equation that describes the change in local diversity over time:

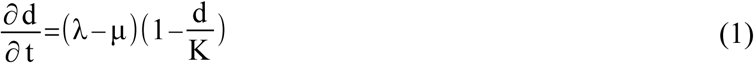

where d is diversity, t is time, λ is the speciation rate, μ is the background extinction rate, and K is the system’s carrying capacity. K varies as a linear function of the net primary production (NPP), ranging between K_min_ and K_max_ corresponding to the 1 and 99 percentiles of the NPP throughout the Phanerozoic.

We model λ according to the following equation:

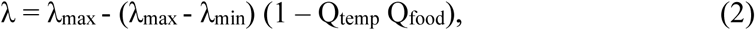

where λ_min_ and λ_max_ are the range of the maximum potential values of λ. Specifically, λ_max_ is the maximum potential speciation rate under optimal environmental conditions. Q_temp_ and Q_food_ are dimensionless factors, ranging from 0 (maximum limitation) to 1 (no limitation), that modulate λ as a function of temperature and NPP or food supply, respectively.

Q_temp_ is calculated using the equation:

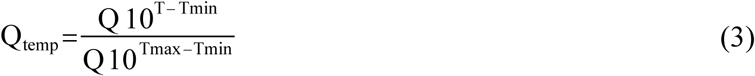

where Q10 is a coefficient that quantifies the sensitivity of λ to temperature, T is the sea surface temperature (°C) at a specific location and time, and T_min_ and T_max_ represent the 1 and 99 percentiles, respectively, of the T frequency distribution within each time interval.

Q_food_ is modelled using a Michaelis-Menten formulation:

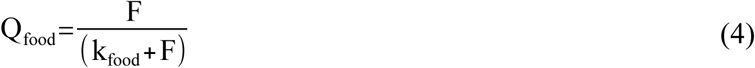

where F (g C m^−2^ yr^−1^) is marine export production, that is, the fraction of net primary production (NPP) that escapes the sunlit surface layer, at a given location and time. The parameter k_food_ (g C m^-2^ yr^-1^) is the value of F at which λ is half of its λ_max_. The same type of Michaelis-Menten parameterization can be used for modelling water availability in terrestrial systems as its effect on λ is similarly dependent on its influence on NPP.

In this study, the background extinction rate (μ) is a fixed value that represents the consistent, low-level rate of diversity loss that occurs even in an ideal environment. It is a fundamental component of the potential net diversification rate (ρ_potential_), defined as:

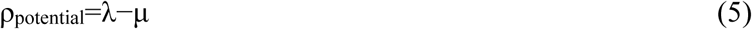

However, this potential rate is not always realized. The realized net diversification rate (ρ_realized_), which is what is actually observed, is constrained by ecological interactions that intensify as d approaches the system’s K:

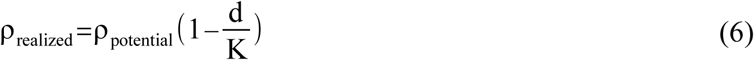

As d nears K, ρ_realized_ declines toward zero, a reduction implicitly caused by a decrease in λ, an increase in μ, or a combination of both (**Fig. 1**). Crucially, K is not a fixed value; it is directly linked to NPP (or food availability). Consequently, a decrease in NPP due to environmental shifts lowers the system’s K. This resource limitation triggers a local extinction to align the value of d with the new, reduced K.

Finally, mass extinctions are simulated by imposing negative net diversification rates. Genus-level marine invertebrate fossil diversity curves are used to determine the percentage of diversity loss, starting time, and duration of each mass extinction (Alroy, 2010; Sepkoski, 2002; Zaffos et al., 2017). To obtain a more robust estimate, independent of biases in individual databases, fossil diversity curves from the various sources are averaged into a single dataset.

The Phyton code for the INDITEK-2.0 model is available at: https://github.com/INDITEK-eco-evo-Model/indicios2.0

### 2.3. Tracking diversification in space and time

Plate tectonics, through the fragmentation and coalescence of continental land masses, profoundly influenced the spatial patterns of biodiversity by creating geographic barriers to species dispersal and spurring provinciality (Pellissier et al., 2018; Valentine et al., 1978; Valentine & Moores, 1970; Zaffos et al., 2017). The shifting of continents and oceans also moved marine and terrestrial habitats across bioclimatic regions, further influencing diversification dynamics. However, previous eco-evolutionary models lack the capability to continuously track the environmental history and biodiversity of habitats as they moved with plates, precluding us from establishing a quantitative, mechanistic link between plate tectonics, global climate evolution, and macroevolutionary dynamics. To mechanistically account for the interplay of plate tectonics and climate on macroevolutionary dynamics, INDITEK implemented a Lagrangian approach (Cermeno et al. 2022). This approach allows the model to track habitats—represented as discrete palaeogeographic points by the plate tectonic model—as they move across Earth’s surface. In doing so, it accounts for the contribution of continuously-evolving environmental conditions and the legacy of past biodiversity changes. A Lagrangian approach contrasts with the simulation of biodiversity ‘snapshots’ (Pohl et al., 2023) and provides a more realistic view of how biodiversity evolved in space and time, capturing the continuous nature of macroevolutionary processes. Currently, integrated modelling frameworks are used to reconstruct Earth’s palaeoenvironmental conditions.

For example, plate tectonic and landscape evolution models provide detailed insights into the planet’s palaeogeographic and physiographic evolution (Kocsis & Scotese, 2021; Merdith et al., 2021; Salles et al., 2023; Scotese & Wright, 2018; Vérard et al., 2015). Likewise, coupled Earth system models and ocean-atmosphere general circulation models reconstruct palaeoclimatic and palaeoenvironmental conditions (Gurung et al., 2024; Mills et al., 2021; Ridgwell et al., 2007; Scotese, 2021). By simulating key environmental variables, such as air/sea temperature, precipitation, and primary production in land and ocean, these models collectively provide the essential context for the unfolding of eco-evolutionary processes (**Fig. 2a**).

**Fig. 2.**
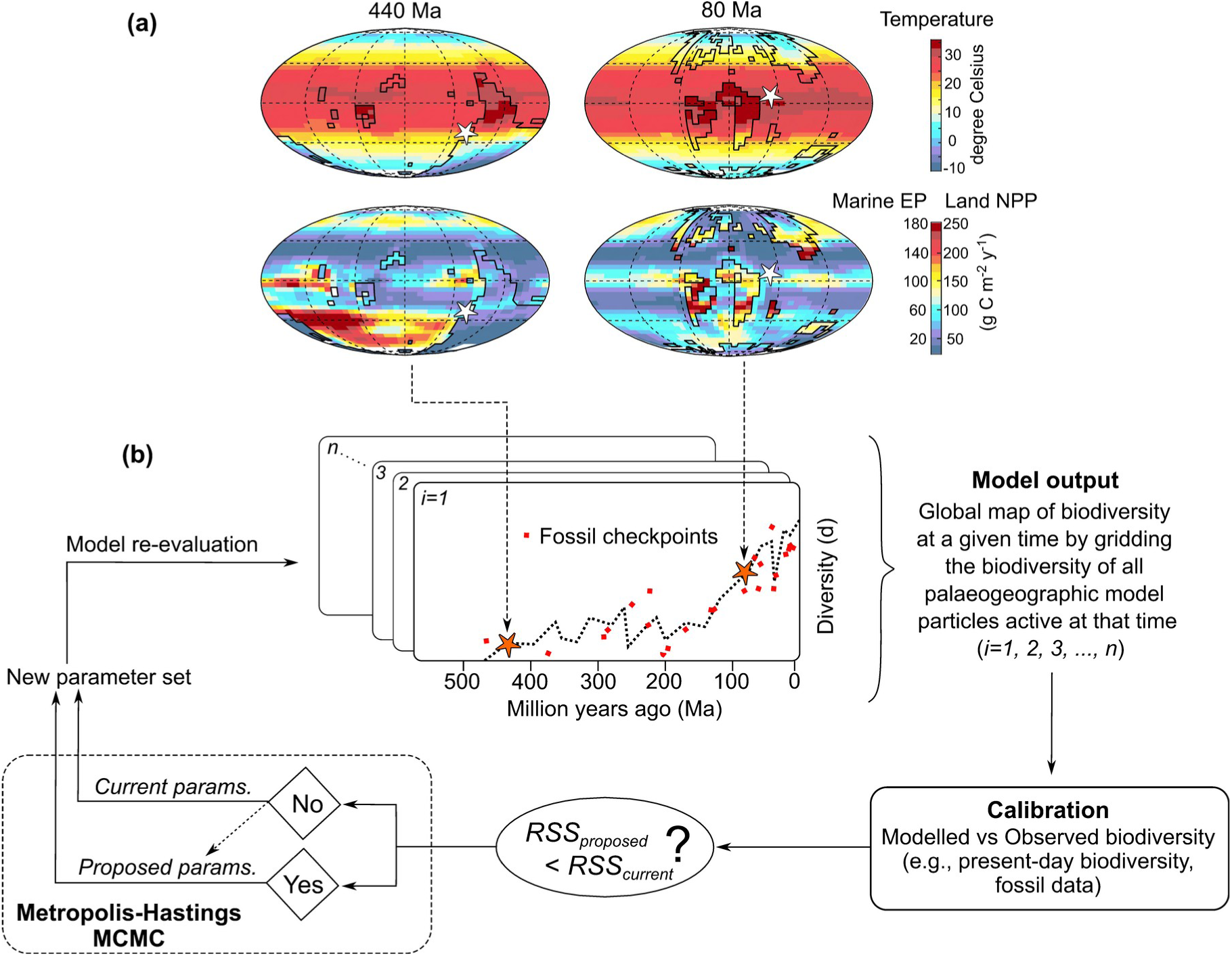
A schematic of the Bayesian inverse modelling framework. **(a)** Global reconstructions of key environmental variables for two distinct time points: 440 Ma (million years ago) and 80 Ma. The top panels show reconstructed air-sea temperatures, while the bottom panels display marine export production (EP) and land net primary production (NPP). The black stepped outlines show the positions of continents. **(b)** Flow diagram of the Bayesian inverse modelling process. The method uses a Metropolis-Hastings Markov chain Monte Carlo (MCMC) algorithm to estimate model parameters by iteratively comparing simulated biodiversity to a calibration dataset. The residual sum of squares (RSS), the likelihood function, quantifies the discrepancy between the modelled and observed biodiversity (or calibration) data, allowing the algorithm to find the best-fitting parameter values.

### 2.4. Bayesian inverse eco-evolutionary modelling

INDITEK-2.0 uses a Bayesian inverse modelling framework with a Metropolis-Hastings Markov chain Monte Carlo (MCMC) algorithm to efficiently explore the parameter space (Hastings, 1970). The MCMC algorithm is used to generate a series of samples from the posterior probability distribution of the model parameters, which allows us to find the most probable parameter values and quantify their uncertainty. In each iteration, the algorithm proposes a new parameter set by randomly perturbing the current set. It then calculates the Residual Sum of Squares (RSS) as a measure of fit to compare the model’s simulated biodiversity and the calibration data (e.g., present-day and fossil biodiversity data) (**Fig. 2b**). New parameter sets are accepted if they reduce the RSS or according to a probabilistic criterion. This criterion allows for the occasional acceptance of less-optimal sets to avoid getting stuck in a local minimum. This iterative process creates a random walk through the parameter space that eventually converges on the posterior distribution, which is the probabilistic solution. After a sufficient “burn-in” period, the collected samples of parameter values approximate this distribution, providing a comprehensive representation of the probability of different parameter values given the observational data and our prior assumptions. This approach ensures that our parameter estimates are not single point values but a range of plausible values, reflecting the uncertainties inherent in the empirical data and the model processes. The model’s parallelized architecture allows multiple independent chains to run simultaneously, which significantly increases the coverage of the sampled parameter space without requiring additional computational time.

### 2.5. Proof-of-concept design

To assess our approach and demonstrate its resolving potential, we performed a synthetic test, or proof-of-concept. First, we selected the target parameter values based on the results of Cermeño et al. (2022) for the continental shelves (**Table 1**). Due to a lack of prior information, we assigned uniform priors with relatively broad range limits to all parameters except for the Q10 parameter; for this parameter, we used a Gaussian distribution, as ample literature for marine invertebrates reports a constrained range of values (1–4), with a higher probability around the mean (1.5–2.5) (Brockington & Clarke, 2001; R. E. Rangel & Sorte, 2022).

**Table 1.** INDITEK-2.0 model parameters: prior distributions and posterior results.

| Parameter | Units | True value | Prior distribution | Posterior Mean | 95% CI |
| --- | --- | --- | --- | --- | --- |
| $K_{\min}$ | #genera | 19 | Uniform (1, 150) | 18.81 | [18.19, 19.6] |
| $K_{\max}$ | #genera | 161 | Uniform (80, 1000) | 162 | [154.63, 170.63] |
| $\lambda_{\min}$ | Myr<sup>-1</sup> | 0.001 | Uniform (0.0001, 0.05) | 0.0019 | [0.0005, 0.0028] |
| $\lambda_{\max}$ | Myr<sup>-1</sup> | 0.035 | Uniform (0.001, ∞) | 0.0349 | [0.0344, 0.0352] |
| $\mu$ | Myr<sup>-1</sup> | 0.001 | n/a | n/a | n/a |
| Q10 | dimensionless | 1.75 | Gaussian (1 – 4) | 1.731 | [1.621, 1.833] |
| $k_{\text{food}}$ | dimensionless | 1.5 | n/a | n/a | n/a |

With these parameter values, we ran the model to generate a synthetic dataset representing present-day biodiversity distributions on a 0.5° x 0.5° spatial grid. In order to mimic observational and theoretical errors and to enable efficient parameter exploration with the Metropolis–Hastings MCMC algorithm, we added a random Gaussian error with a standard deviation of 2 genera to the synthetic dataset. We then applied our inversion approach to this dataset. Six Markov chains were used to explore the 5-dimensional parameter space and to sample the posterior probability distribution. Each chain was initialized within a range of ±20% of the reference parameter values, with starting points selected using a Latin hypercube sampling strategy to ensure broad coverage of the parameter space. At each iteration, one forward simulation was performed, and the resulting output was compared to the synthetic dataset. Model fit was quantified using the residual sum of squares (RSS), calculated as the squared difference between the simulated and synthetic data, normalized by the variance of the added noise.

## 3. Results

The MCMC chains for all five parameters: K_max_, K_min_, λ_max_, λ_min_, and Q10 converged toward a stable mean over the 2000 interative simulations after successful exploration of the posterior distribution (**Fig. 3**). The rate of convergence varied among parameters. K_max_, K_min_, and λ_max_ showed relatively rapid convergence, with chains stabilizing within 500 to 750 iterations. In contrast, Q10 and particularly λ_min_ showed slower convergence, with some chains for λ_min_ taking nearly 1500 iterations to mix around the original, true value. The convergence of all chains confirms the stability and reliability of the parameter estimates.

**Fig. 3.**
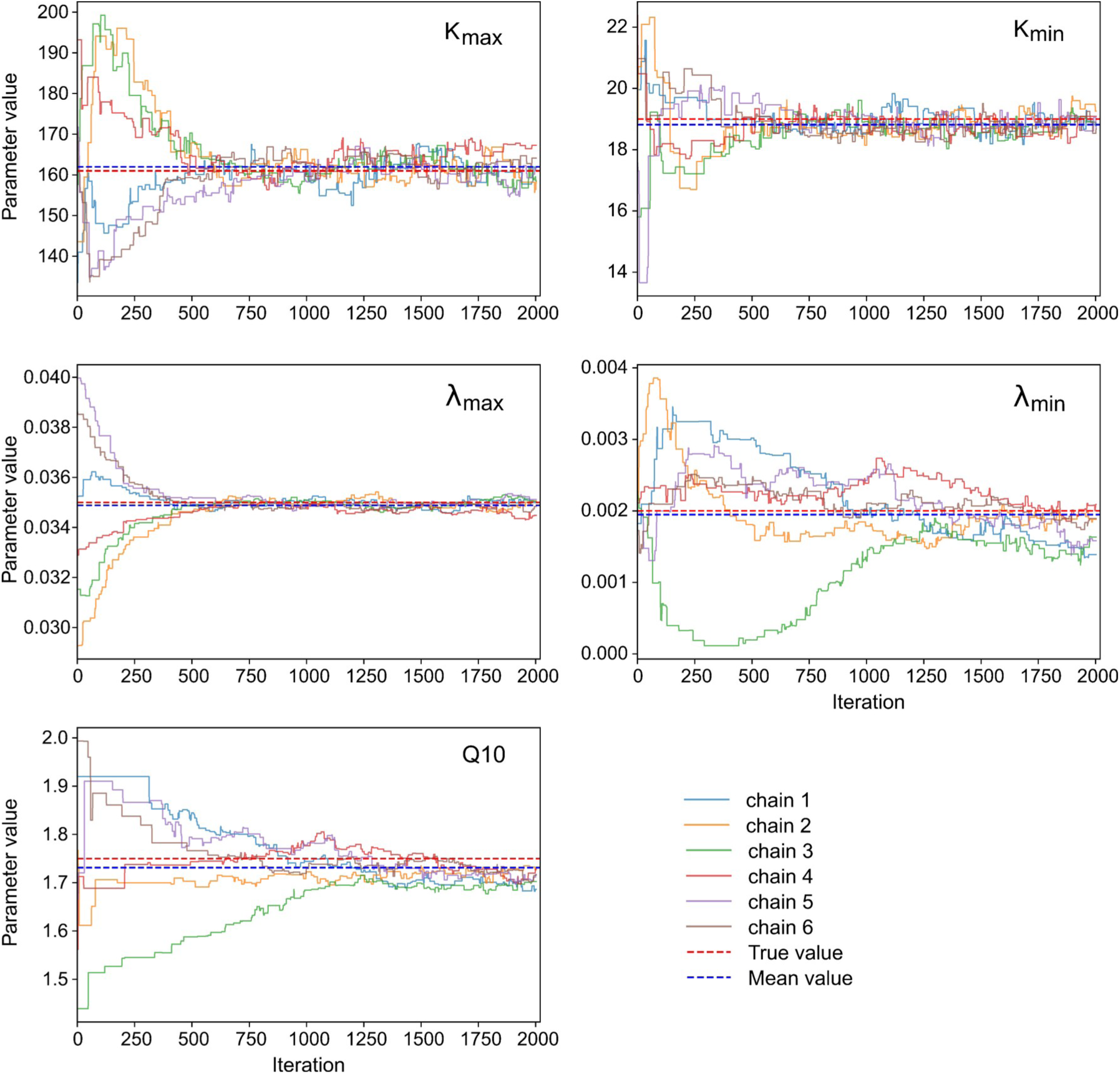
Markov chain trajectories for parameter inference. Evolution of five model parameters (K_max_, K_min_, λ_max_, λ_min_, and Q10) over 2000 iterations for each of the six independent Markov chains (labeled 1 through 6). The horizontal dashed red line indicates the true value for each parameter, while the horizontal dashed blue line indicates the mean value of the posterior distribution after the burn-in period (from iteration 200 onward).

Our analysis further confirms the accuracy of the model, as the true parameter values were always contained within the 65% credible interval provided by the posterior solution (**Table 1**, **Fig. 4**). This demonstrates that the MCMC method not only found a stable solution but also accurately estimated the range of plausible parameter values, reliably capturing the true underlying values within this interval. The two-dimensional joint posterior distributions correctly identified the relationships between parameter pairs (**Fig. 4**). The bivariate correlations among the model parameters offered key insights into the structure of the parameter space and potential interdependencies. A strong positive correlation was observed between λ_min_ and Q10, suggesting that the MCMC algorithm compensated for a lower λ_min_ by exploring higher values for Q10. Conversely, K_max_ and K_min_, the upper and lower bounds of the carrying capacity showed, respectively, a negative and positive correlation with λ_max_. This might indicate a trade-off in how the model fits the synthetic biodiversity data. Specifically, to explain biodiversity patterns with a very high λ_max_, the model must constrain K_max_ to prevent a runaway increase that would overshoot the data. In contrast, a high λ_max_ is positively linked to K_min_, indicating that rapid diversity growth is necessary to explain a high baseline level of biodiversity. In essence, the model balances a high-speed diversification process against both K_max_ and K_min_ to reproduce the synthetic biodiversity data. Other parameter pairs showed weak or negligible correlation, as indicated by their generally more circular joint posterior distributions, suggesting that the MCMC algorithm explored these parameters with a relatively high degree of independence (**Fig. 4**).

**Fig. 4.**
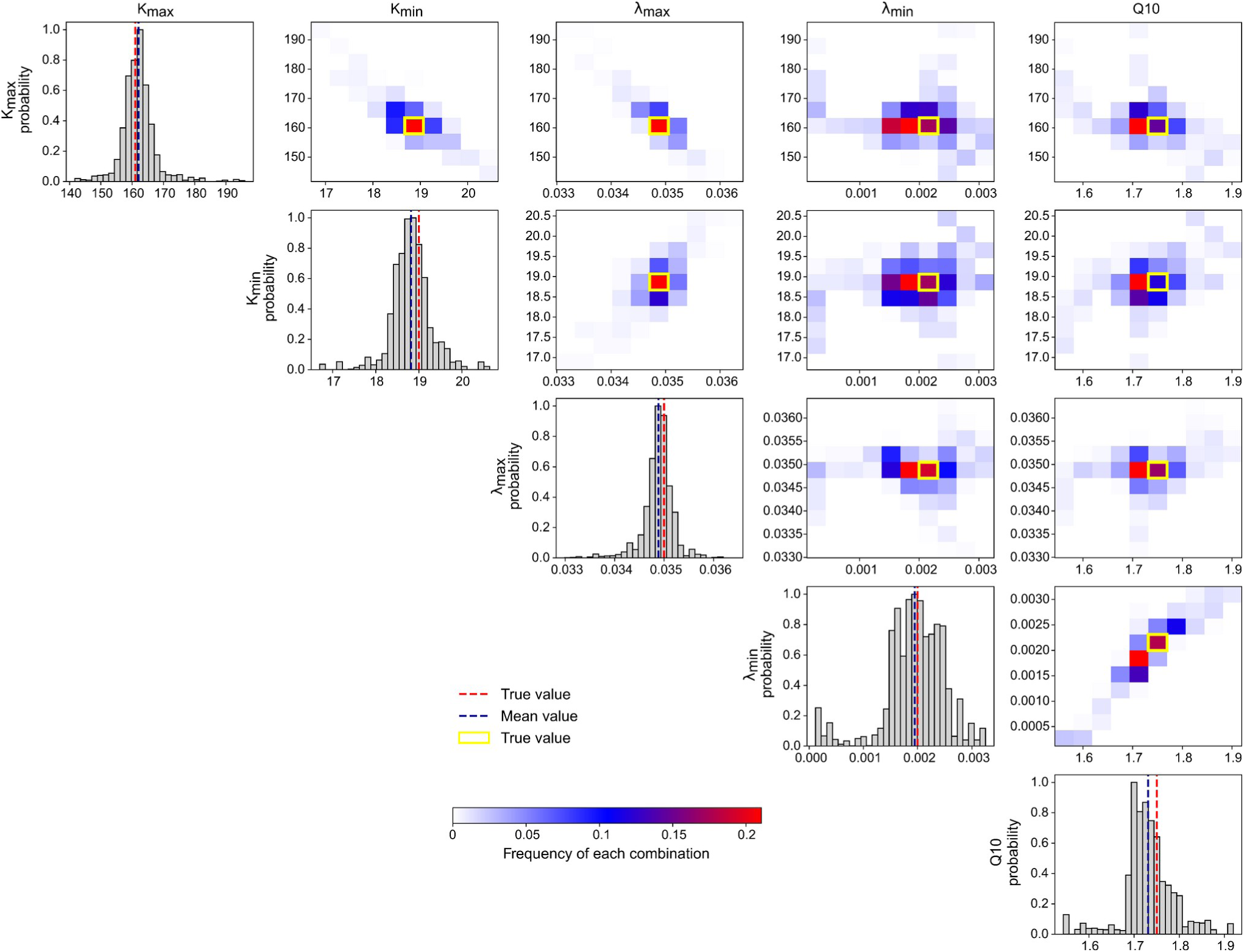
Validation of the INDITEK-2.0’s Bayesian inverse modelling framework. The diagonal panels show the one-dimensional posterior distributions for each of the five inferred parameters (K_max_, K_min_, λ_max_, λ_min_, and Q10), represented as histograms of all pooled MCMC samples after the burn-in period. The dashed red line indicates the true value for each parameter, while the dashed blue line shows the mean value of the posterior distribution after the burn-in. The off-diagonal panels display the joint posterior distributions and correlations between parameter pairs. The colour gradient denotes the density of MCMC samples. The red square included on each heatmap represents the true value used to generate the synthetic biodiversity data. The location of this red square relative to the densest region of the heatmap is a key indicator of the framework’s success in recovering the original parameter values.

We found a strong and regionally balanced match between the true, synthetic biodiversity data and the estimated biodiversity data derived from the mean of the probabilistic solutions (**Fig. 5**), indicating the model’s high performance in parameter estimation. The minimal differences between the two biodiversity distribution maps do not detract from the model’s overall accuracy.

**Fig. 5.**
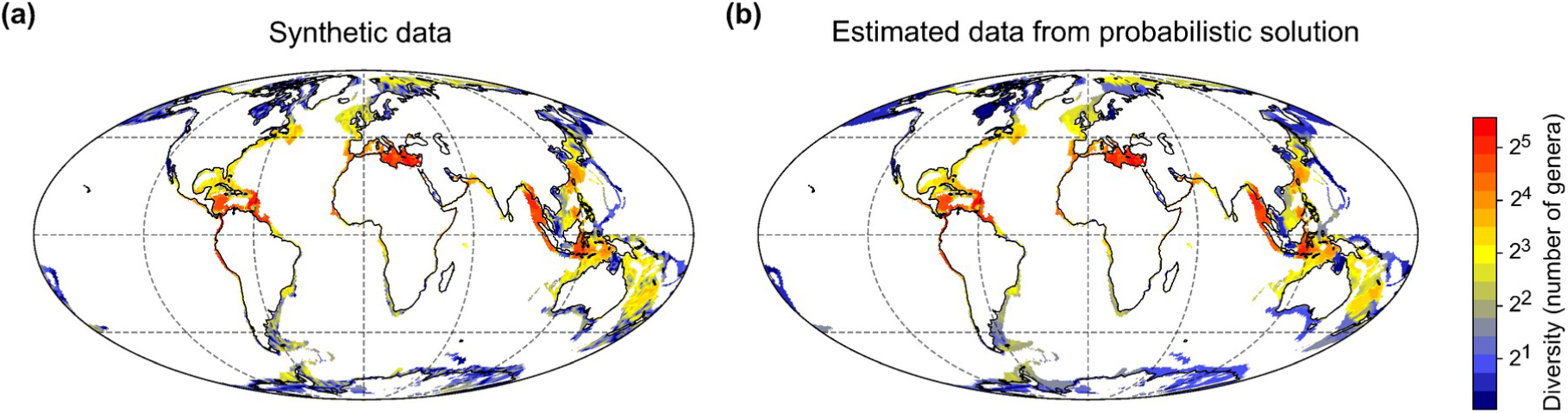
Marine invertebrate biodiversity data. **(a)** Global distribution of marine invertebrate diversity generated using the INDITEK model with predefined (true) parameter values. **(b)** Global distribution of marine invertebrate diversity derived from parameters estimated using the INDITEK-2.0’s Bayesian inversion method.

Our approach generated an ensemble of plausible biodiversity histories rather than a single definitive one. Each history, with its assigned probability, allowed us to explicitly quantify the uncertainty in our reconstructions. To illustrate this uncertainty, we tracked the biodiversity of three representative modern regions; the Indo-Pacific, Caribbean, and Mediterranean submerged habitats, throughout the simulations (**Fig. 6**). These regions, now among the most striking marine biodiversity hotspots worldwide, began to develop from the Late Triassic onwards, likely due to the relatively high long-term environmental stability that prevailed during the Mesozoic and Cenozoic eras (Cermeño et al. 2022). Our analysis was designed to test the Bayesian inverse modelling approach, so the variability in our reconstructions was relatively low and mostly stemmed from the error we deliberately introduced to the synthetic data (**Fig. 6**). We anticipate a higher variability in biodiversity reconstructions when model simulations are calibrated with true, field observational data. The analysis of this variability will be crucial in future studies to pinpoint which macroevolutionary patterns and drivers are most robustly supported by real-world evidence.

**Fig. 6.**
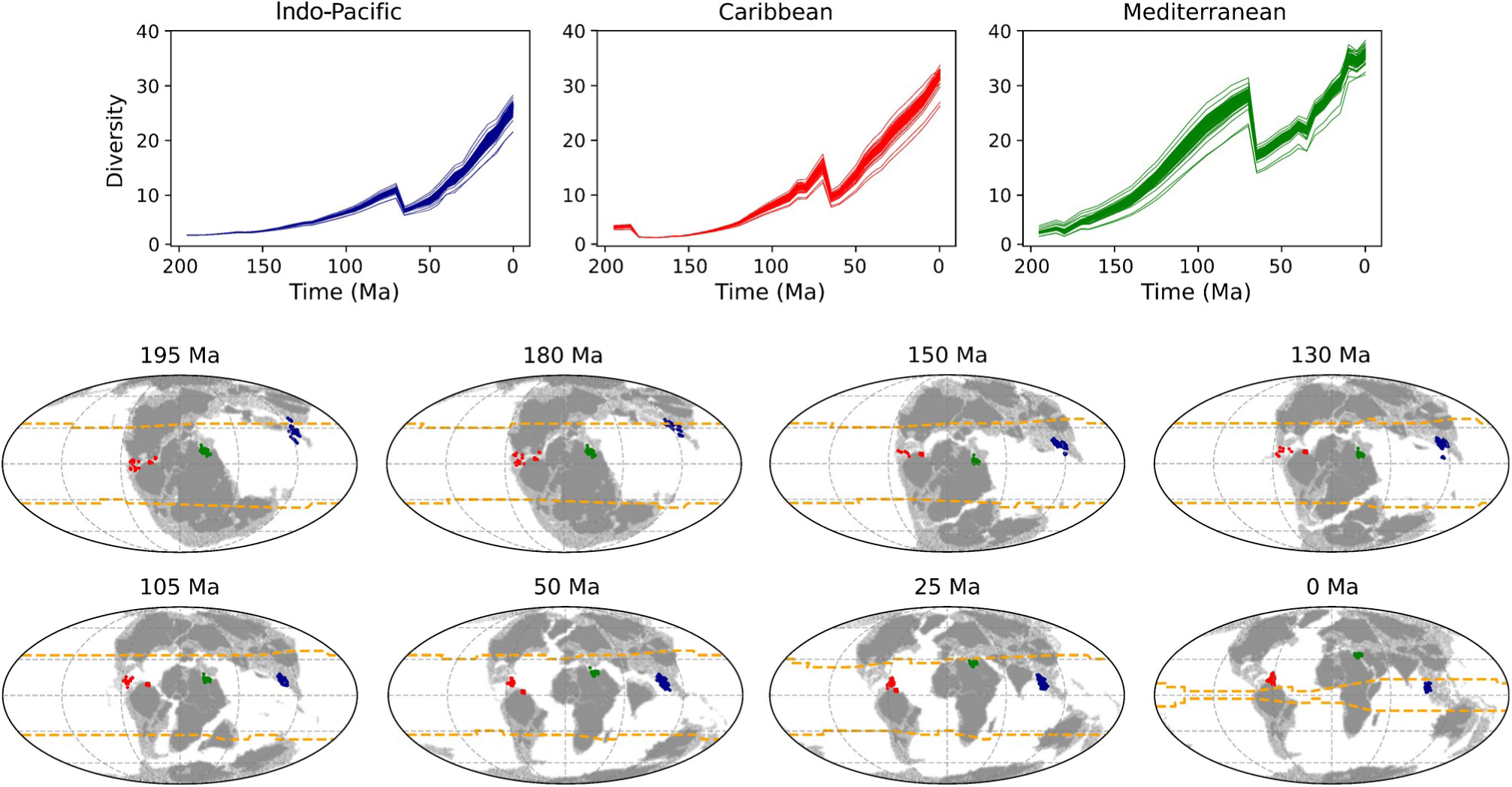
Macroevolutionary history of marine biodiversity hotspots. (upper panels) Temporal evolution of marine invertebrate diversity (median number of genera across all tracked palaeogeographic points) from the Late Triassic (∼200 Ma) to the present day (0 Ma) for the Indo-Pacific, Caribbean, and Mediterranean regions. Each panel displays multiple histories, corresponding to the different probabilistic solutions from the model. **(lower panels)** Geographic positions of the tracked palaeogeagraphic points for each biodiversity hotspot (colour dots) at specific time intervals. Each map shows the continental configuration (dark grey) and continental shelves (light grey) at the indicated time. The orange dashed lines represent the approximate positions of the 25°C isolines (isotherms).

## 4. Caveats and future directions

Our mechanistic modelling tool provides a virtual window into a profoundly complex evolutionary history. Evolutionary processes are shaped by many factors, many of which are beyond the current possibilities of the model. Recognizing these limitations is crucial to adequately interpreting the model’s results and devising future implementations.

i. The extreme environmental changes associated with mass extinctions pose a significant challenge for modelling due to the involvement of multiple stressors, often interacting in nonlinear ways. For example, the Ordovician mass extinction (ca. 445 million years ago) is thought to have been initiated by global glaciation, which was followed by a rapid warming period. These climate fluctuations led to rapid changes in sea level and ocean anoxia with dramatic consequences for marine life (Pohl et al., 2023). Similarly, the end-Permian mass extinction is thought to have been caused by massive volcanic eruptions, leading to severe climatic shifts, ocean acidification, and widespread anoxia (Hülse et al., 2021). Modelling mass extinctions requires sophisticated parameterizations to capture nonlinear dynamics and feedback among Earth’s evolving systems (geosphere, atmosphere, hydrosphere, and biosphere). These systems are intrinsically interconnected, with changes in one domain often leading to cascading effects in others (Hull, 2015; Junium et al., 2022; Schobben et al., 2019). The difficulty in implementing parameterizations that capture these nonlinear dynamics and feedback loops in palaeo-Earth system models currently poses significant challenges in recreating the environmental scenarios responsible for mass extinctions.
ii. As discussed in ***Section 2.1. Parameterizing evolutionary rates***, adaptation allows species to persist in changing environments. However, rapid environmental changes can outpace the rate of evolutionary adaptation, preventing species from developing the necessary traits to cope with new conditions. This reasoning posits a direct linkage between the rate of environmental change and the rate of species extinction (Kaiho, 2022; Song et al., 2021). Implementing this parameterization is currently limited by lack of palaeoenvironmental reconstructions with sufficient spatial and temporal resolution. Our current model uses environmental inputs from reconstructions with a temporal resolution of 30 time slices each encompassing 10–20 million-year and a spatial resolution of 36 x 36 (latitude, longitude) equal area grid. Extinction events, however, are not uniform across vast geographic areas or over long timescales; instead, they are often localized and triggered by relatively rapid environmental shifts. As a result, existing coarse-resolution reconstructions may smooth out these crucial, fine-scale variations, resulting in an inaccurate estimation of the extinction rate (μ). Therefore, the successful application of this parameterization is contingent on the availability of high-resolution environmental reconstructions that can accurately capture the nature of extinction dynamics. Current efforts are being made to increase these resolutions to every million years and 72 x 72 equal area grid.
iii. Extinction selectivity is influenced by countless biological, ecological, and environmental variables, often interacting unpredictably. The ecological niches, life history traits, and geographic distributions of species significantly influence their resilience or vulnerability to environmental perturbations (Cole & Hopkins, 2021; Payne et al., 2016; Pinsky et al., 2019). These species-specific attributes cannot be implemented into a model operating above the species level. However, it is feasible to implement extinction selectivity at a higher taxonomic level by adjusting specific parameter values associated to a class or phylum known for their overall sensitivity or resistance to particular stressors. For example, assigning different values to ectotherms and endotherms for the thermal dependence of speciation rate can reflect their differential responses to temperature changes. Insects’ short generation times, and, in many cases, ability to go dormant can confer resistance to environmental stressors. Their evolutionary success could be modelled and parameterized by, for example, lowering their sensitivity to mass extinctions.
iv. The emergence of innovative evolutionary traits and co-evolutionary relationships has been pivotal in creating novel ecological opportunities and driving major increases in biodiversity (Benton et al., 2015; Nitecki, 1990; Vermeij, 2017). At or below the species level, mechanistic eco-evolutionary models can be designed to simulate how new traits emerge and evolve. These models can include mechanisms for trait evolution (e.g., trait diffusion), niche construction, and ecological interactions, allowing them to capture the interplay between organisms and their environments that drives speciation (Hagen, 2023). To account for the effect of evolutionary innovations and co-evolutionary relationships on macroevolutionary dynamics, future work could involve using fossil data and molecular phylogenetic analyses to identify specific time points that mark key innovations or the start of co-evolutionary histories. We could then estimate model parameters for the periods before and after these critical time points, allowing the eco-evolutionary model to represent discrete, biologically-justified shifts in evolutionary dynamics and carrying capacities. For example, evolutionary innovation can expand the ecospace and thus the amount of diversity supported per unit resource supply, which is parameterized as an increase in K_min_ and K_max_.

## 5. Conclusions

Our study presents INDITEK-2.0, a cost-efficient mechanistic eco-evolutionary model designed to explore the deep-time history of biodiversity. Initially tested on synthetic communities of marine invertebrates, the model is extensible to other marine and terrestrial animal groups, microorganisms and plants, provided that 1) comprehensive and reliable calibration datasets are available, and 2) the model’s parameter-dependent processes realistically represent their evolutionary dynamics.

INDITEK-2.0 is built on two innovative approaches that allow us to trace the ancient origins of today’s biodiversity patterns. First, unlike traditional models that use a fixed geographic grid, INDITEK-2.0 employs a Lagrangian approach. This method tracks specific parcels of the Earth’s surface as they move over millions of years due to plate tectonics. This dynamic tracking is essential for accurately linking modern regions of interest, such as biodiversity hotspots, to their ancient, non-stationary geographic locations and environmental conditions. Second, INDITEK-2.0 utilizes a Bayesian inverse modelling framework to probabilistically estimate model parameters, thereby generating hundreds to thousands of plausible biodiversity histories, each with a statistically quantified probability. By integrating these two methods, INDITEK-2.0 offers a powerful and novel way to investigate the link between Earth’s surface dynamics and biodiversity history.

INDITEK-2.0 allows us to target a specific taxonomic group and reconstruct its biodiversity history by iteratively refining model outputs based on comparisons with both present-day biodiversity and fossil data. While probabilistic methods have long been used to detect diversification rate shifts from time-calibrated molecular phylogenies (Bouckaert et al., 2019; Morlon et al., 2024; Rabosky, 2014; Stadler, 2011), applying these methods to spatially-explicit mechanistic eco-evolutionary models offers the ability to glean novel insights into the mechanisms driving these shifts. This is a key distinction. By interrogating the model, we can directly investigate how, for example, continental drift and global climate evolution influenced macroevolutionary dynamics, providing an explicit and realistic representation of the environmental forces that shaped biodiversity over millions of years. Ultimately, integrating mechanistic eco-evolutionary models with probabilistic parameter estimation provides a robust, uncertainty-aware framework for reconstructing biodiversity’s lost history. This approach represents a significant step toward uncovering the origins of the astonishing biodiversity we see today.

## Acknowledgements

This work was funded by research grant PID2023-152076NB-I00 from the Spanish government. We thank Lewis Jones and Thiago Rangel for comments on an early version of the manuscript. T.B. acknowledges the support from the Spanish Ministry of Science and Innovation ATR2023-143517/AEI/10.13039/501100011033. A.P. acknowledges the support of the French *Agence Nationale de la Recherche* (ANR) under reference ANR-22-CE01-0003 (project ECO-BOOST). Calculations were performed using HPC resources from DNUM CCUB (*Centre de Calcul de l’Université de Bourgogne*) and the SMART HPC infrastructure at the Institut de Ciències del Mar. P.C., C.G.C, G.H.G. and T.B. acknowledge the ‘Severo Ochoa Centre of Excellence’ accreditation (CEX2019-000928-S).

## Supporting information

### Earth system model simulations

This section describes the methods used to simulate the Late Ordovician and Late Cretaceous palaeoenvironmental conditions shown in the figure 2 of the manuscript. To date, no numerical model offers a coupled representation of climate-carbon-cycle equilibrium, primary productivity on land and marine biogeochemistry in a spatially-resolved, 3D ocean – which together constitute the boundary conditions needed for Phanerozoic-scale mechanistic eco-evolutionary model simulations. This limitation arises from the contrasting time scales at which the underlying mechanisms occur.

Carbon cycle equilibrium, on the one hand, results from the balance between (silicate, carbonate and kerogen) weathering on land and (carbonate and organic carbon) burial in the ocean, and operates at the million-year time scale. Oceanic circulation and biogeochemistry, on the other hand, reach equilibrium much faster, in a few 10s of kilo-years. As a result of this major difference in the time constants of the different components of the climate system, the long-term carbon cycle and the ocean-atmosphere system have traditionally been simulated using very different types of models. Long-term carbon cycle models can be run for 100s of million years. They can be used to simulate the evolution of the atmospheric concentration in greenhouse gases (CO_2_ in particular) and oxygen, and changes in oceanic nutrient inventories (Mills et al., 2021). Their rapid integration time relies on parameterizations and spatial abstraction – generally representing the ocean and atmosphere as a single box (hence value). Earth system models, on the other hand, offer a spatially-resolved simulation of climate and ocean biogeochemistry, but their resulting computational cost usually does not permit to run them for more than a few 10s of kyrs (Pohl et al., 2022).

To overcome this limitation, we here propose to couple a long-term carbon cycle model (SCION) (Mills et al., 2021) with an Earth system model of intermediate complexity (cGENIE) (Ridgwell et al., 2007). In detail, we propose to first simulate palaeoclimatic conditions using an ocean-atmosphere general circulation model (e.g., FOAM) (Jacob, 1997). Palaeoclimatic conditions simulated at various atmospheric CO_2_ levels can then be used to calculate the long-term climate-carbon-cycle equilibrium using SCION, together with primary productivity on land using the vegetation module FLORA (Gurung et al., 2022, 2024). Atmospheric CO_2_ and O_2_ concentrations and oceanic nutrient (phosphate) inventories simulated in SCION can be used as boundary conditions in cGENIE, the latter model permitting in turn to simulate ocean temperatures and biogeochemistry (including marine export production) in a 3D ocean. During all steps, the same continental reconstructions are used in every model. The final result is a self-consistent set of palaeoenvironmental conditions both on land and in the ocean – notably temperatures and primary productivity that are two key drivers of speciation and extinction rates.

